# Analysis of patient derived xenograft studies in Oncology drug development: impact on design and interpretation of future studies

**DOI:** 10.1101/579136

**Authors:** Jake Dickinson, Marcel de Matas, Paul A Dickinson, Hitesh Mistry

## Abstract

**Background:** Preclinical Oncology drug development is heavily reliant on xenograft studies to assess the anti-tumour effect of new compounds. Patient derived xenograft (PDX) have become popular as they may better represent the clinical disease, however variability is greater than in cell-line derived xenografts. The typical approach of analysing these studies involves performing an un-paired t-test on the mean tumour volumes between the treated and control group at the end of the study. This approach ignores the time-series and may result in false conclusions, especially when considering the increased variability of PDX studies.

**Aim:** To test the hypothesis that a model-based analysis provides increased power than analysis of final day volumes and to provide insights into more efficient PDX study designs.

**Methods:** Data was extracted from tumour xenograft time-series data from a large publicly available PDX drug treatment database released by Novartis. For all 2-arm studies the percent tumour growth inhibition (TGI) at two time-points, day 10 and day 14 was calculated. For each study, the effect of treatment was calculated using an un-paired t-test and also a model-based analysis using the likelihood ratio-test. In addition a simulation study was also performed to assess the difference in power between the two data-analysis approaches for different levels of TGI for PDX or standard cell-line derived xenografts (CDX).

**Results:** The model-based analysis had greater statistical power than the un-paired t-test approach within the PDX data-set. The model-based approach was able to detect TGI values as low as 25 percent whereas the un-paired t-test approach required at least 50 percent TGI. These findings were confirmed within the simulation study performed which also highlighted that CDX studies require less animals than PDX studies which show the equivalent level of TGI.

**Conclusion:** The analysis of 59 2-arm PDX studies highlighted that taking a model-based approach gave increased statistical power over simply performing an un-paired t-test on the final study day. Importantly the model-based approach was able to detect smaller size of effect compared to the un-paired t-test approach is which maybe common of such studies. These findings were confirmed within simulated studies which also highlighted the same sample size used for CDX studies would lead to inadequately powered PDX studies. Application of a model-based analysis should allow studies to use less animals and run experiments for a shorter period thus providing effective insight into compound anti-tumour activity

## Introduction

Preclinical Oncology drug development is heavily reliant on xenograft studies to assess the anti-tumour effect of new compounds [1]. These studies are often quite short in duration, on the order of 1 to 2 weeks, and represent the first opportunity, during development, to assess how the kinetics of drug disposition affects the kinetics of tumour growth. The xenograft study starts by grafting a human cell-culture into the flank of an immunocompromised mouse. Digital callipers are then used to measure the length and width or the length, width and height to calculate tumour volume at regular time-intervals. Once the grafted tumour has reached a certain pre-specified volume each xenograft is randomised into one of the treatment arms or the untreated arm (control arm within the study). Thus, across all study arms the volume at randomisation is comparable. The treatment effect is then calculated by measuring the difference in mean tumour volumes between the treated and control group at the end of the study. This metric is typically referred to as the Tumour Growth Inhibition (TGI) value and is calculated as follows:

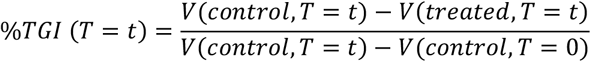

where *V(treated, T=t)* and *V(control, T=t)* are the mean volumes of the treated and control group at time *T=t*, the end of the study, and *V(control, T=0)* is the mean volume of the control group at the time of randomisation. To assess whether the volumes at time *t=T* are significantly different between the control and treated arms of the study an unpaired t-test is performed and the resultant p-value is usually reported together with the TGI value.

The above approach to analysis of tumour growth inhibition data clearly ignores the time-series that is generated up until the TGI value is recorded. Furthermore, the TGI value can become biased if mice have dropped out at a time-point before the TGI value is calculated. This is common in the control arm due to the volume exceeding a pre-defined animal welfare limit. Thus, performing an un-paired t-test on the final day of the study, results in an under-prediction of the mean control volume and hence the efficacy of the treatment is underestimated. This issue however can be resolved by calculating TGI at an earlier time-point where no drop-outs exist or a joint longitudinal drop-out model could also be used [2].

An alternative to performing an un-paired t-test on the final study day is to perform a model-based regression analysis. The key advantage of performing a regression analysis over the approach discussed is that all data points in the time series are used. This will lead to an increase in statistical power, hence reduce the number of animals used and thus reduce the cost of a xenograft study.

A previous study, by Hather et al. [3], has shown that a model-based regression approach is likely to improve the power of xenograft studies over doing an un-paired t-test. However, the Hather et al. study did not consider the application of such an approach to patient-derived xenografts, which are known to have higher variance than the standard cell-line xenograft and are becoming more popular within preclinical development. Furthermore, the approach by Hather et al. did not consider the use of mixed-effects/hierarchical modelling approach which is likely to further increase the statistical power of such studies. In this study we build on the work by Hather et al. by assessing the increase in power obtained by using a model-based mixed-effects regression analysis over the typical unpaired t-test analysis of final volumes for a large open patient derived xenograft database [4]. In addition we also analysed a traditional standard cell-line xenograft study for comparison to the patient-derived xenograft analysis and perform a brief simulation study highlighting the merits of a model-based mixed-effects regression analysis.

## Methods

### Real data study

#### Xenograft data

Data from a study by Gao et al. [4] which analysed 1000 Patient Derived Xenograft (PDX) models across 59 treatments and controls was collected. The data was then grouped creating 59 two arm studies involving a control and treatment arm. This data was then truncated to either 10 or 14 days, to mimic the typical length of a xenograft study, for analysis.

#### Empirical Approach: Unpaired T-test

On the final day of the study, day 10 or 14, an unpaired t-test between the control and treated volumes was conducted with a p-value and %TGI reported.

#### Model-based Approach: Likelihood ratio-test

The time-series from the animals used within the empirical approach were used for the following model-based analysis. We first converted tumour volume to radius using:

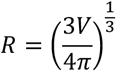

where V is the volume of the tumour and R the radius.

Next, we fitted the following model [5, 6], to the combined two arm data-set:

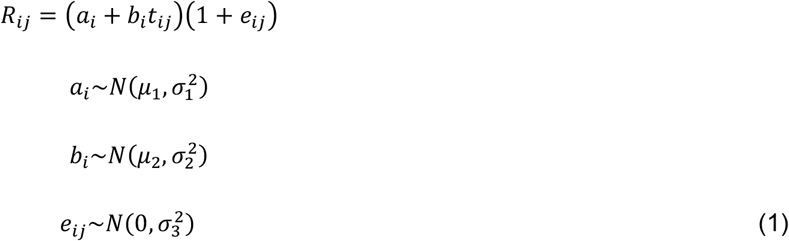

Where *R_ij_* is the observed radius of xenograft *i* at time *j, a_i_* is the value of the radius at time 0, time of randomisation, for xenograft *i, b_i_* is the rate of growth of the radius for xenograft *i, t_ij_* is the time-point for xenograft *i* at time *j* at which the observation was recorded and *e_ij_* is the residual error for xenograft *i* at time *j*. Note, that a proportional error model was used as the variability in tumour size grew over time i.e. we have heteroscedastic variance (see appendix for more details). We then modified the distribution of *b_i_* and introduced a population treatment effect parameter *c* to account for difference between control and treated growth rates in the following way

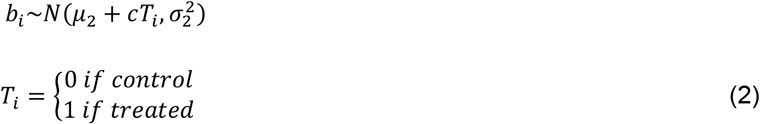

and re-fitted the model to the data. Since the models considered are nested the likelihood ratio-test was used to assess if adding a treatment effect improved model fit to the data over a model with no treatment effect. All parameter estimation was conducted using the *saemix* library in R.

### Simulated data study: power calculation

A 14 day 2-arm simulation study, using model (1) stated above, was conducted in the following way. All parameter values except for the treatment effect *c* were based on fitting model (1) to all control data up to day 14. The treatment effect parameter c was chosen to give either a 50% or 100% TGI at 14 days. The sample size for each arm of the simulated study was set at 5, 8,10, 12 and 15. When simulating the tumour volumes we assumed that the measurements were taken on days 0, 3, 7, 10 and 14. These values were chosen to mimic the Gao et al. study. In addition to using the Gao et al. study to generate model simulation parameters we also utilised a standard cell-line (CDX) study by Knutson et al. [7] which was taken from datadryad.org [8].

For each simulated study, control and treatment arm, the empirical and model-based approaches were applied with p-values recorded. This was done a 1000 times for a given simulation setup with the proportion of p-values <0.05 recorded and visualised.

## Results

### Real data study

The results of the 10 day 2-arm, treatment versus control, study analysis of the PDX data can be seen in Figure 1. We found that 91% of studies had a p-value <0.05 when using a model-based analysis whereas only 31% had a p-value <0.05 using an empirical analysis. Furthermore, we found that the trend for increased power was study independent as the model-based approach is more powerful regardless of the PDX model used, see Figure 2. That is for all studies the p-value obtained using the model-based approach were lower than the empirical approach.

**Figure 1.**
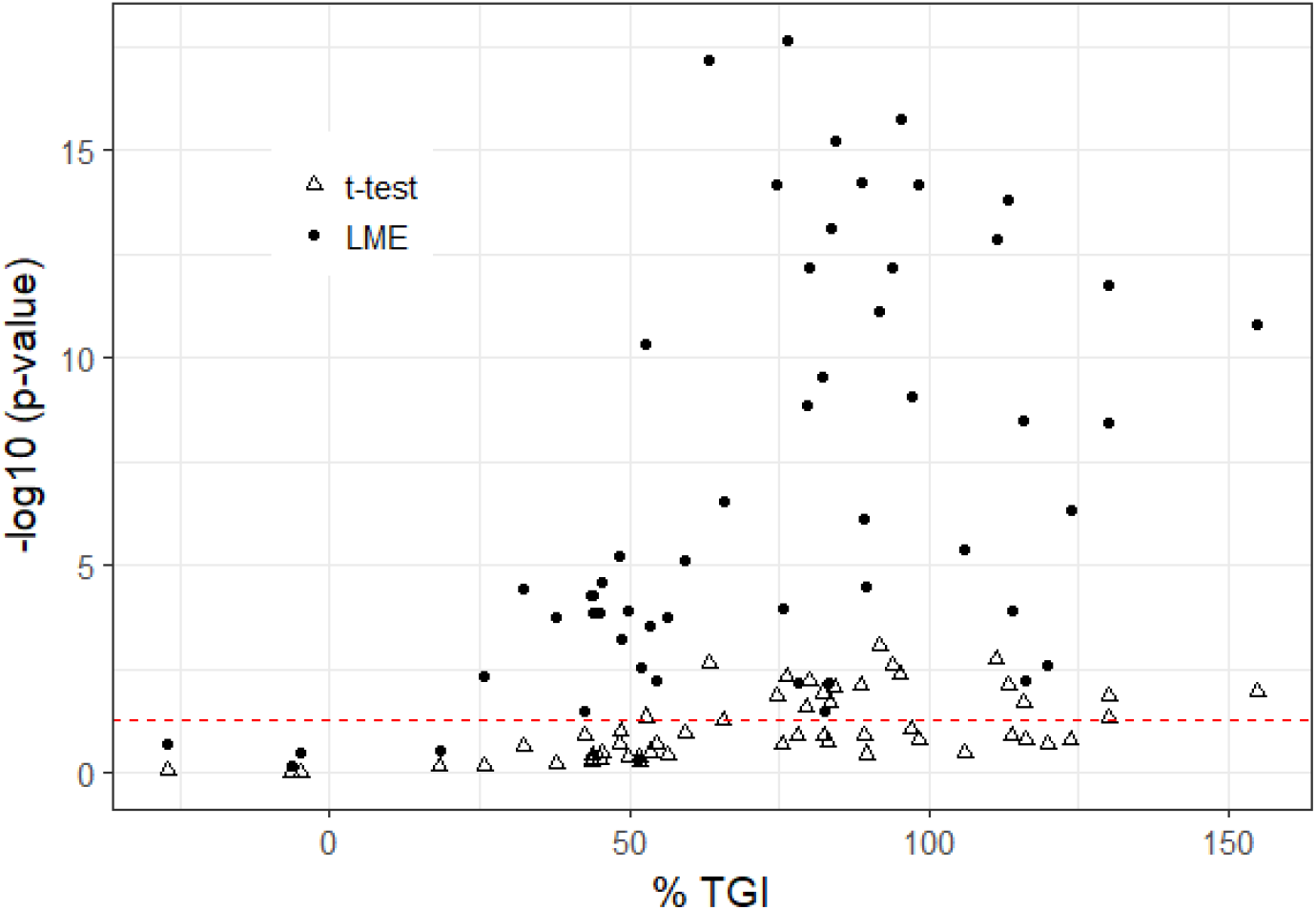
Statistical power of a t-test (Δ) and LRT (likelihood ratio-test) (•). Log transformed p-value plotted against % TGI with a red dashed line indicating the p-value threshold (0.05) after 10 days of treatment.

**Figure 2.**
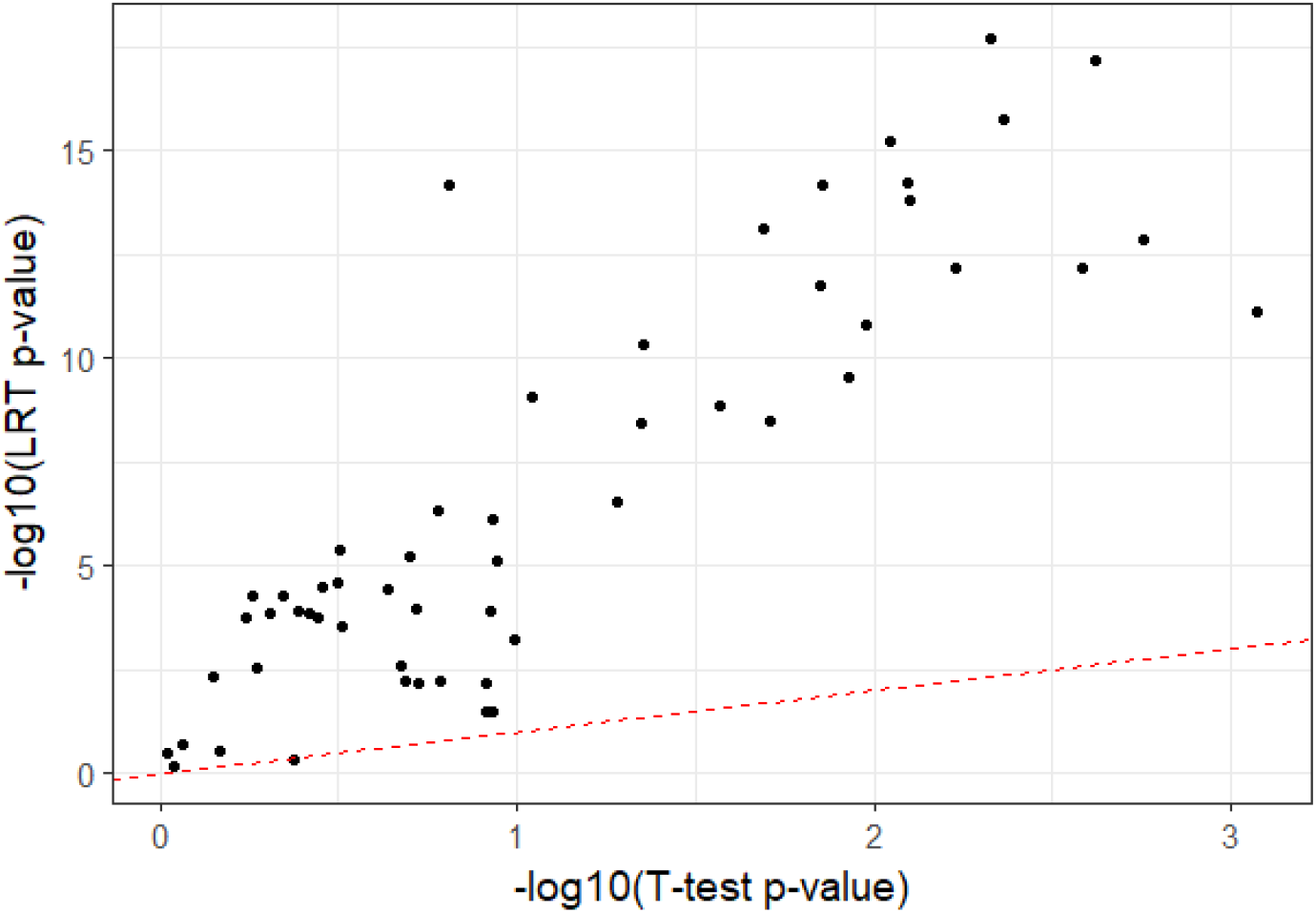
Log transformed LRT (likelihood ratio-test) p-values plotted against log transformed t-test p-values with the line of unity plotted as a dashed redline after 10 days of treatment.

A similar set of results was found when performing a 14 day analysis; see Figure 3 and Figure 4. That is the model-based approach had increased power over the empirical approach. For the 14 day analysis we found that 95% of the studies had a p-value <0.05 via the model-based approach whereas only 44% had a p-value <0.05 using the empirical approach.

**Figure 3.**
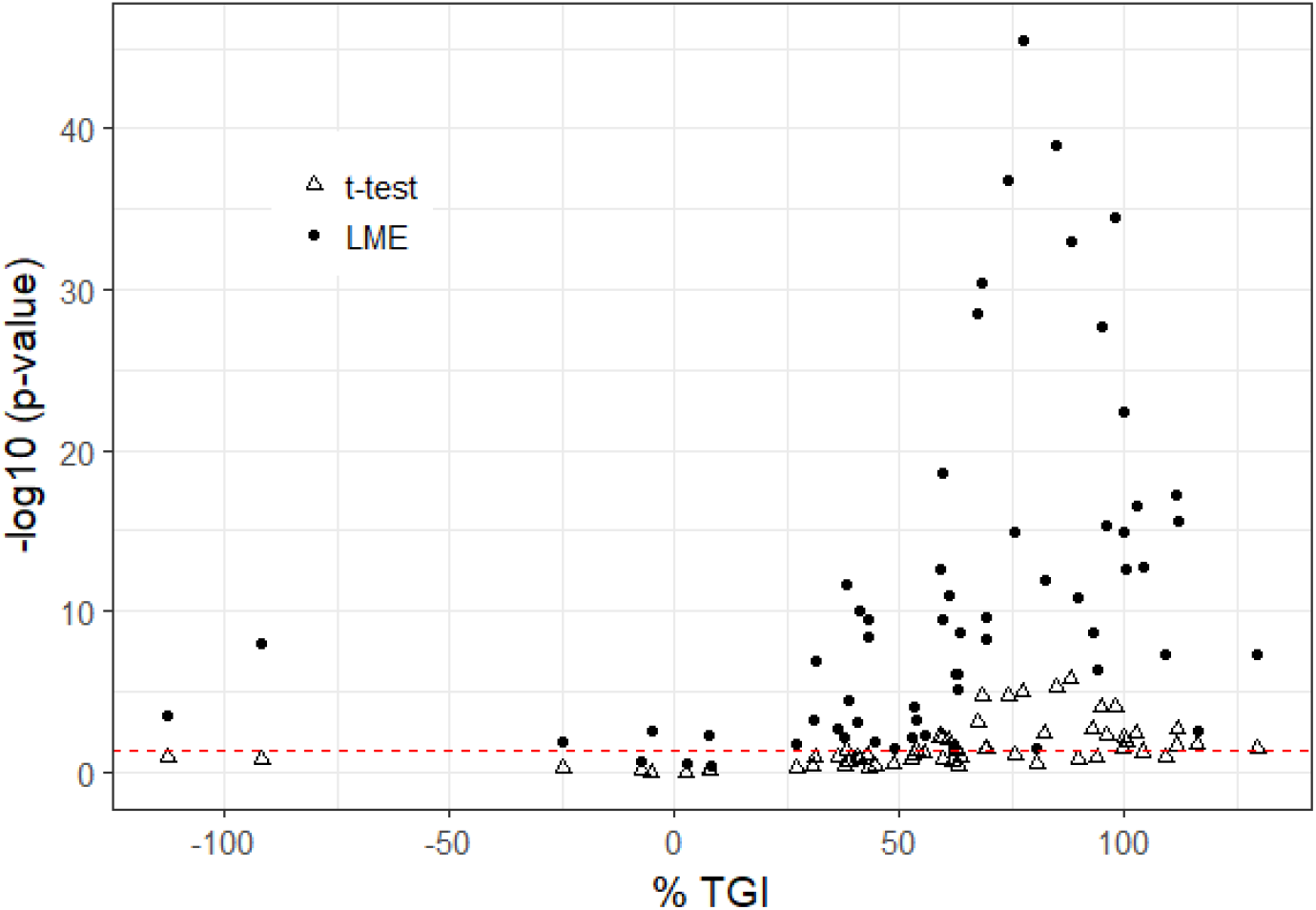
Statistical power of a t-test (Δ) and LME model (•). Log transformed p-value plotted against % TGI with a red dashed line indicating the p-value threshold (0.05) after 14 days of treatment.

**Figure 4.**
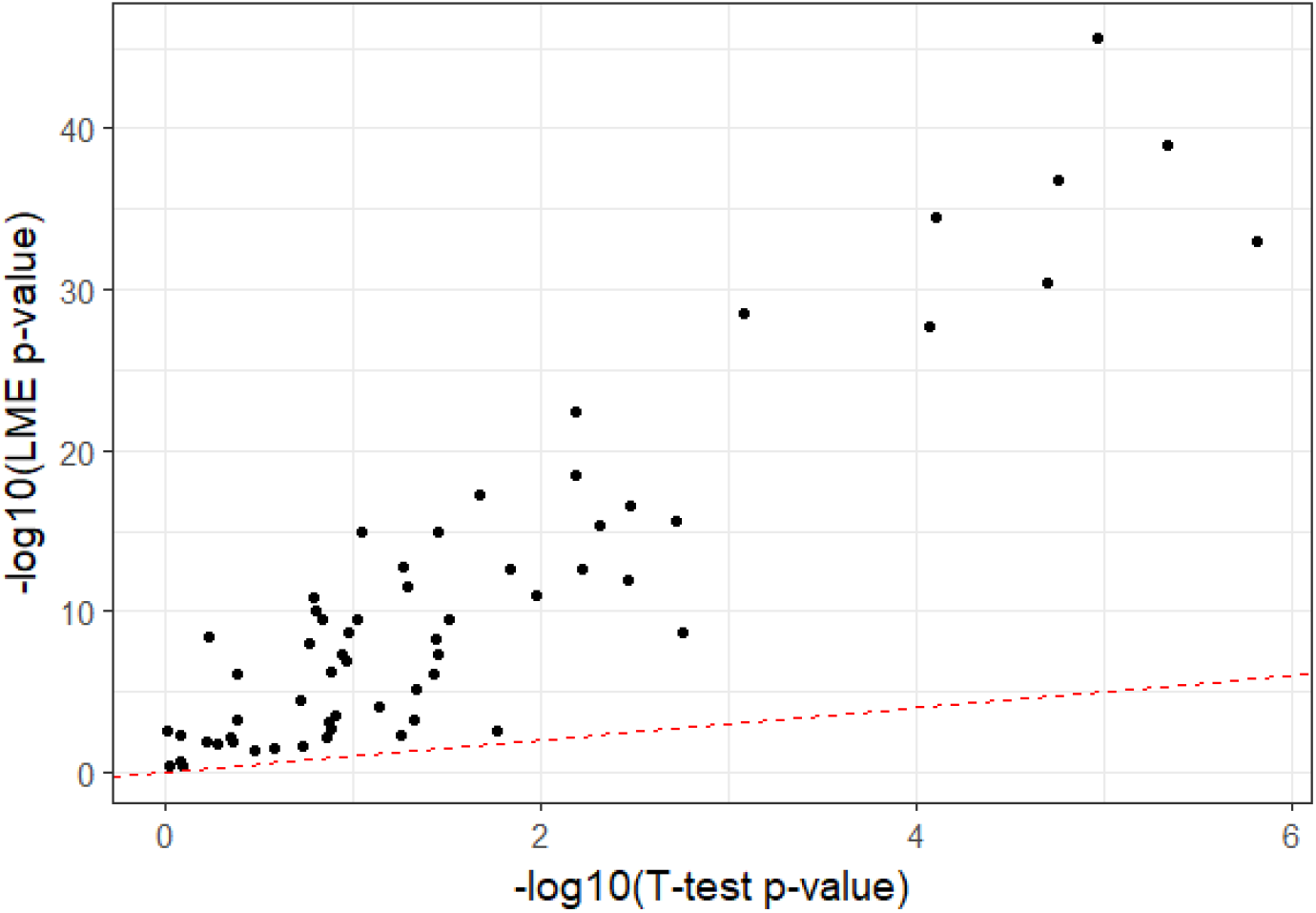
Log transformed LRT (likelihood ratio-test) p-values plotted against log transformed t-test p-values with the line of unity plotted as a dashed redline after 14 days of treatment.

Overall these results show that within the PDX database using the model-based approach leads to significantly more power than the empirical approach of analysis.

To further illustrate the greater power of a model-based approach the cumulative fraction of significant (p<0.05) results was plotted against TGI, see **Error! Reference source not found**.. The Figure shows that the model-based approach was able to detect TGI differences as low as 50% unlike the empirical approach. Additionally, for both methods, a 14 day study has increased power over a 10 day study.

### Simulated data study

The results of the simulation study using parameters derived from analysing the controls of the PDX study can be seen in Table 1. It shows that the power of the model-based approach is greater than that using the empirical approach. We can see that the power does decrease as we decrease the number of mice, as expected. A further simulation power analysis was done using parameters derived from a CDX study, details on the parameter values can be found in supplementary material. The result of this analysis in comparison to the PDX analysis can be seen in Table 2. The results show that if the variability is low and a 100% TGI is sought then the power of using an empirical versus model-based approach is similar. However, in the other scenarios explored we found that the model-based approach has greater power over the empirical approach. The table further highlights the difference in power between CDX and PDX studies; this is due to the increased variability in time-series seen in PDXs, see supplementary material for parameter estimates.

**Table 1.** Results of the simulated PDX power analysis from the empirical and model-based approach.

| Percentage of p-values <0.05 |  |  |  |
| --- | --- | --- | --- |
|  |  | Empirical | Model-Based |
| TGI (%) | N | Day 14 | Day 14 |
| 100 | 15 | 77.5 | 89.2 |
|  | 12 | 67.9 | 77.1 |
|  | 10 | 59.7 | 73.8 |
|  | 8 | 43.9 | 63.1 |
|  | 5 | 24.2 | 44.2 |
| 50 | 15 | 13.7 | 24.3 |
|  | 12 | 9.3 | 12.4 |
|  | 10 | 8.9 | 13.3 |
|  | 8 | 6.1 | 10 |
|  | 5 | 3.7 | 8.1 |

TGI – tumour growth inhibition

**Table 2.** Comparison of PDX and CDX power analyses.

| Percentage of p-values <0.05 |  |  |  |  |  |  |
| --- | --- | --- | --- | --- | --- | --- |
|  |  |  | Empirical |  | Model-Based |  |
| TGI (%) | N | Day | CDX | PDX | CDX | PDX |
| 100 | 10 | 14 | 99.6 | 59.7 | 100.0 | 73.8 |
| 50 |  |  | 55.9 | 8.9 | 78.5 | 13.3 |

TGI – tumour growth inhibition; CV – coefficient of variation; CDX – cell-line derived xenograft; PDX patient derived xenograft

## Discussion

It is well known that cell-line derived xenografts that have been established over the decades poorly mimic human disease. Thus there has been a long-standing interest in developing new animal models that better mimic patient tumours and their microenvironments. One approach that has been gaining favour in recent years is the use of patient derived xenografts. These models show more variability in their time-series and treatment response than their cell-line derived counterparts mainly due to the increased heterogeneity within the sample used. Given the increased cost of using patient derived versus cell-line derived xenografts more importance should be placed on how these studies are analysed than is presently done.

In this study we explored the typical empirical based analysis methods using the final volumes to model-based approaches that use the whole time-series across 59 2 arm trials taken from a publicly available patient derived xenograft database. The empirical approach consisted of applying an unpaired t-test to the final volumes of the control versus treated arms of a study. The model-based approach, however, involved using a parametric model to describe the time-series and the likelihood ratio-test to assess if including a treatment effect parameter improved model fit. Thus the model-based approach used all the data whereas the empirical based approach did not.

The results showed that the model-based approach had more statistical power than the empirical based approach. This is consistent with larger studies involving cell-line derived xenografts that have been previously reported [3]. We found that the model-based approach could identify tumour growth inhibition values as low as 25% whereas >50% tumour growth inhibition is required for the empirical based method to detect a difference. Given that we would expect modest tumour growth inhibition with patient derived xenografts versus cell-line derived xenografts due to increased heterogeneity, this highlights the importance of using a model-based approach for such analyses.

A second key result was that due to lower variability in controls between cell-line versus patient derived xenografts, no difference in statistical power was found when considering 100% tumour growth inhibition. This was not the case when moving to patient derived xenografts, where an increase from 60% to 74% power was observed using a model-based over empirical based approach.

In summary the results have shown that detecting modest TGI values such as 50% a model-based approach will lead to increased power for both CDX and PDX studies. Wong et al. [9] have shown that clinical exposure targets that relate to at least 60% TGI in cell-line derived xenograft studies are required to see clinical efficacy. Thus, based on the power analysis conducted here a model-based analysis would be the most appropriate approach to detect such modest efficacy.

Unlike previous studies conducted within this field we chose to use the same end-point, percent tumour growth inhibition, when comparing analysis methods. In doing so we hope that experimental scientists can see that the way the outcomes are reported need not change only the method of analysis. We hope this will encourage experimental scientists to explore model-based xenograft analysis approaches as we enter the age of more sophisticated animal models in cancer.

## Supplementary Information

### Analysis of PDX controls

We found that using a proportional error model gave a better model fit to the control data truncated at 14 days than using an additive model (−2xlog-likleihood – proportional error = 668; −2xlog-likleihood – additive error = 750). The final model parameters for the control fit can be seen in Table 3, these values were used for the PDX simulation study. A posterior (visual) predictive check can be seen in Figure 5.

**Table 3.** Parameter estimates of the final control PDX model.

| a (mm) |  | b (mm/day) |  | e |
| --- | --- | --- | --- | --- |
| Mean | Variance | Mean | Variance | Variance |
| (%RSE) | (%RSE) | (%RSE) | (%RSE) | (%RSE) |
| 3.83 | 0.06 | 0.10 | 0.0050 | 0.04 |

RSE – relative standard error;

**Figure 5.**
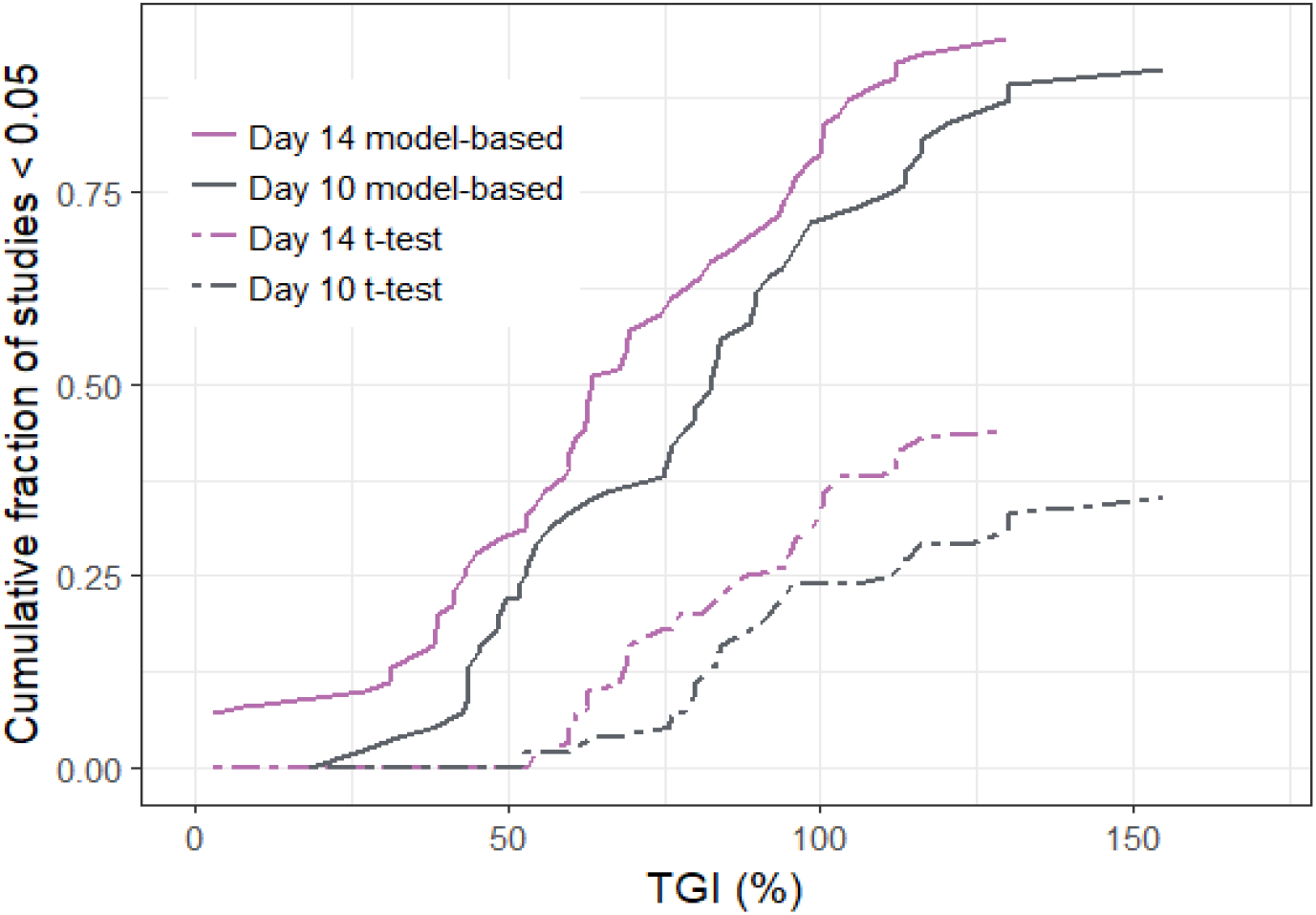
Cumulative fraction of significant studies vs tumour growth inhibition (TGI)

**Figure 5a.**
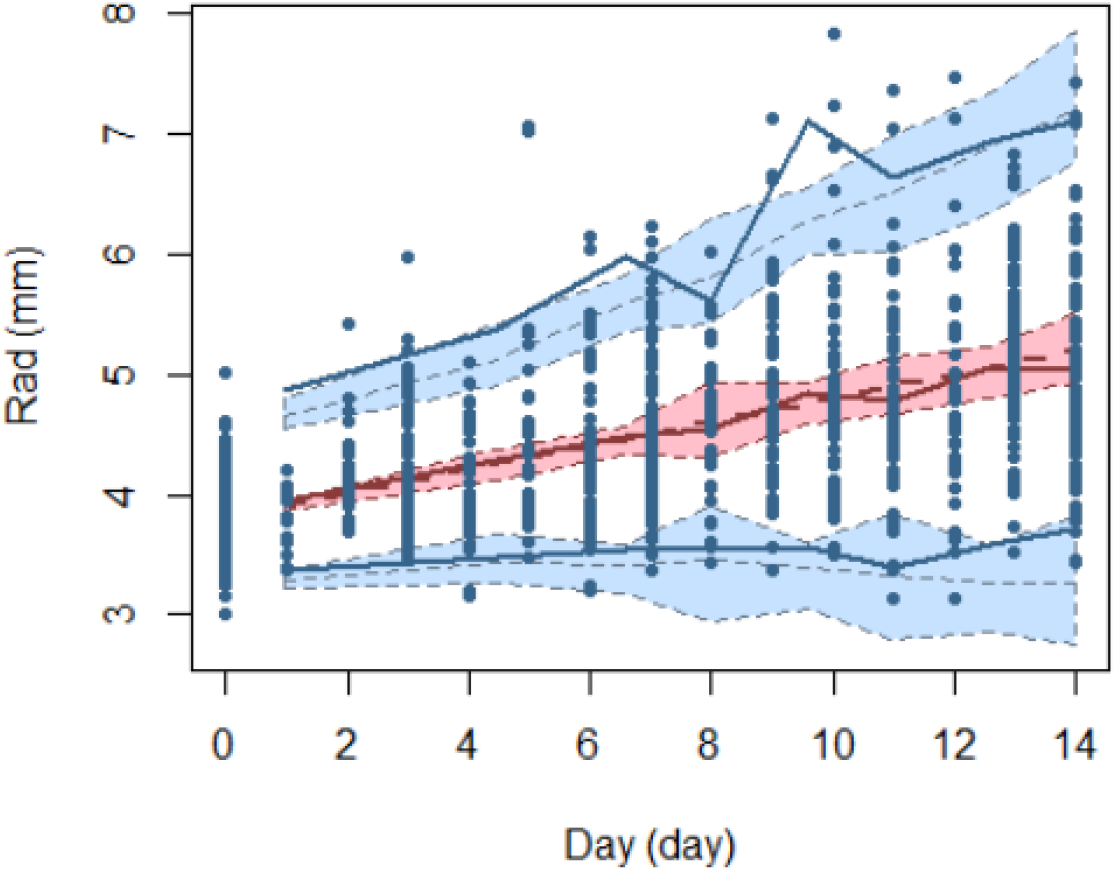
Visual predictive check of the tumour growth model with proportional residual error model.

Solid dots are the observed values, solid red-line is the observed median over time, solid blue lines are the observed 2.5^th^ and 97.5^th^ percentiles over time, the pink shaded region is the 95% confidence interval for the median and the blue shaded regions are the 95% confidence intervals for the 2.5^th^ and 97.5^th^ percentile both derived by simulation.

### Analysis of CDX controls

The final model parameters to the control data, using a proportional error model, in the cell-line derived xenograft study can be seen in supplementary Table 4, these values were used in the CDX simulation study. A posterior (visual) predictive check can be seen in Figure 6.

**Figure 6.**
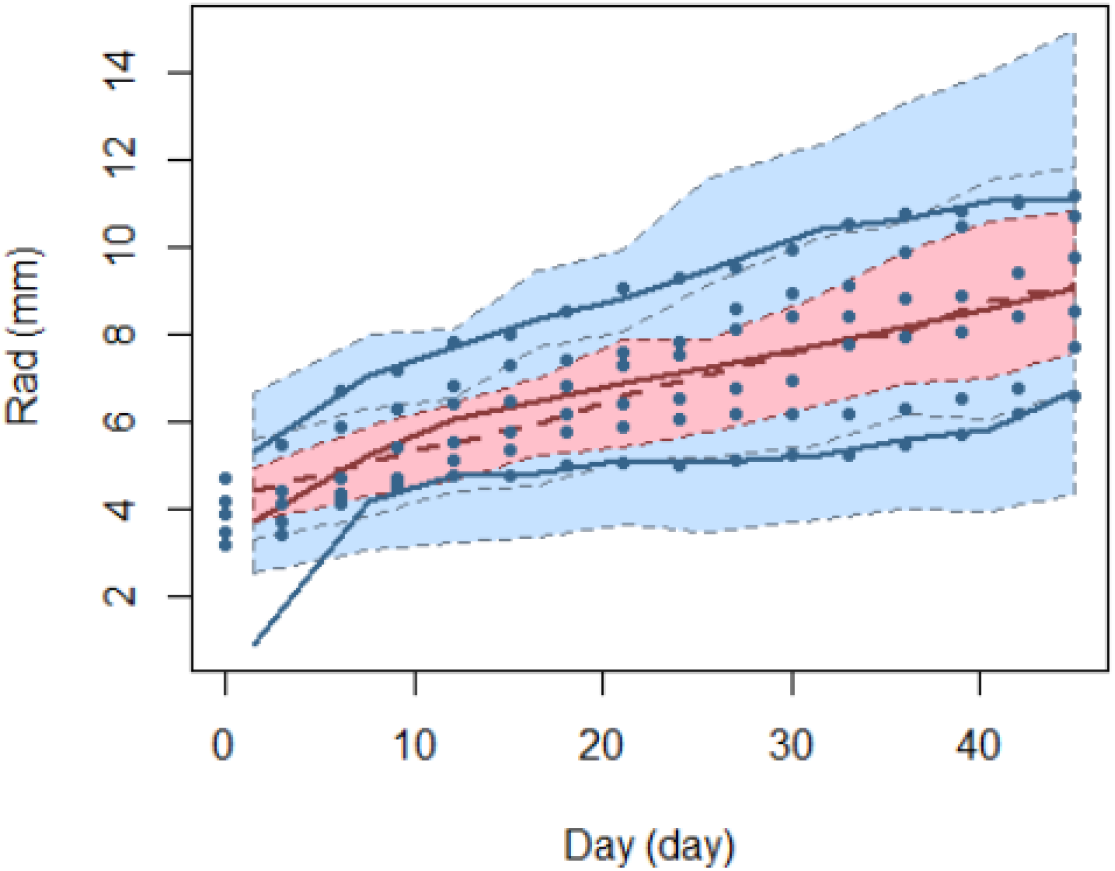
Visual predictive check of the tumour growth model with proportional residual error model.

**Table 4.**
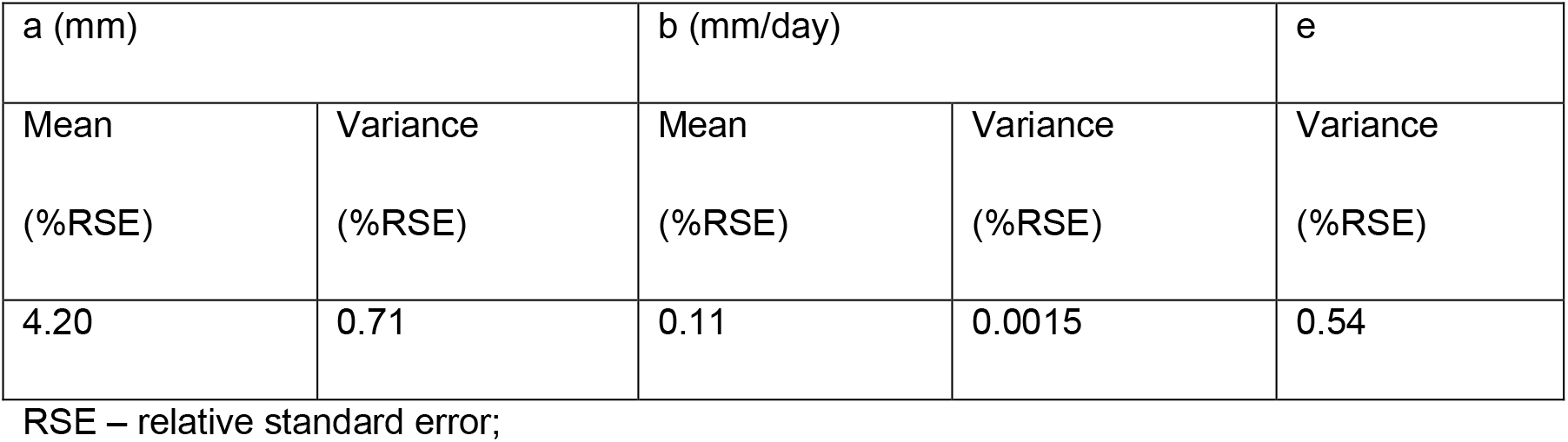
Parameter estimates of the final control CDX model.

| a (mm) |  | b (mm/day) |  | e |
| --- | --- | --- | --- | --- |
| Mean | Variance | Mean | Variance | Variance |
| (%RSE) | (%RSE) | (%RSE) | (%RSE) | (%RSE) |
| 4.20 | 0.71 | 0.11 | 0.0015 | 0.54 |

RSE – relative standard error;

Solid dots are the observed values, solid red-line is the observed median over time, solid blue lines are the observed 2.5^th^ and 97.5^th^ percentiles over time, the pink shaded region is the 95% confidence interval for the median and the blue shaded regions are the 95% confidence intervals for the 2.5^th^ and 97.5^th^ percentile both derived by simulation.

